# Defining Functional Correction Thresholds in Primary Ciliary Dyskinesia for Effective Gene Therapies

**DOI:** 10.64898/2026.02.19.706855

**Authors:** Beck E. Fitzpatrick, Brett J. Wineinger, Jason E. Babcock, Erik J. Quiroz, Emily C. Liu, Lalit K. Gautam, Alejandro A. Pezzulo, Thomas O. Moninger, David K. Meyerholz, Douglas B. Hornick, Amy L. Ryan

## Abstract

**Rationale:** Primary ciliary dyskinesia (PCD) is an inherited disorder characterized by defective motile cilia and impaired mucociliary clearance. Mutations in CCDC40 disrupt axonemal organization, resulting in dyskinetic or immotile cilia. While emerging therapies may restore function in only a subset of cells, the functional consequences of mixed populations of mutant and healthy cilia are not well understood.

**Objectives:** To determine how defined mixtures of CCDC40-deficient and wild-type ciliated cells influence mucociliary transport.

**Methods:** Human bronchial epithelial cells were combined at varying ratios of CCDC40-deficient and wild-type cells and differentiated to model heterogeneous epithelia. High-speed video microscopy and particle-tracking algorithms were used to assess ciliary motion and mucus transport and combined with electron microscopy to evaluate cilia ultrastructure.

**Measurements and Main Results:** Airway differentiation was largely preserved with marked ultrastructural defects observed in CCDC40 cells, including multiple central centrioles, absent inner dynein arms, and basal body misorientation. Ciliated surface coverage decreased, and goblet coverage increased with higher mutant representation. Mucociliary transport declined nonlinearly, with speeds dropping from ∼56 µm/s (100% WT) to ∼9 µm/s in 100% mutant cultures. Clearance-per-beat and flow coordination decreased sharply with rising mutant burden. Modeling revealed that transport efficiency was equivalent to that recorded in ex vivo human tissues and plateaued when ∼75% of the ciliated population was WT.

**Conclusions:** Together, these findings define a PCD-specific functional correction threshold and show that effective therapy must overcome the disruptive biomechanical and cellular influences of mutant epithelial cells, providing a quantitative benchmark to guide gene-therapy design and clinical translation.

## INTRODUCTION

Mucociliary clearance (MCC) is a fundamental defense mechanism of the respiratory tract, reliant on the coordinated beating of motile cilia across a continuous epithelium. This highly integrated, multicellular process ensures the directional transport of mucus and entrapped pathogens out of the airway. Mutations that affect motile cilia cause primary ciliary dyskinesia (PCD) which represents a spectrum of motile cilia abnormalities broadly associated with chronic bronchitis, sinusitis, and atelectasis: cilia may be absent, reduced in number, missing essential sub-components that make up the normal axonemal and basal body architecture, or functionally dis-coordinated^1,2^. For example, mutations in cyclin-O (CCNO), cause a severe form of the disease that disrupts cilia extension^3-7^, and whereas mutations in the dynein chains, such as dynein heavy chain 5 (DNAH5), lead to static cilia^8-10^. Defective ciliary function and the resulting deficiency in MCC, lead to recurrent respiratory tract infections and progressive lung damage. In most instances of PCD, defects in ciliary length, waveform, and beat kinematics are evident^11-14^. Causative mutations have yet to be identified in ∼30% of diagnosed PCD subjects and it is plausible that complex heterozygosity with allelic mutations in two different PCD genes could lead to mild symptomatic disease^15-20^. Mutations in coiled-coil domain-containing 39 and 40 (CCDC39/40), genes critical for the assembly of inner dynein arms and ciliary axonemal organization, result in immotile or severely dyskinetic cilia. These are associated with a particularly severe clinical phenotype^17,21-24^, suggesting that small differences in ciliary defects can manifest in substantially different clinical outcomes.

We recently established a quantitative map of the key luminal cell types, ciliated and secretory cells, in both human and rat airways, revealing significant species-specific differences including much higher levels of ciliation in human compared to rodent large airways^25^. Building on this we demonstrated that mucociliary clearance capacity is tightly coupled to the density and spatial organization of ciliated cells: tissues with higher ciliated cell abundance and more continuous ciliary coverage exhibited markedly improved mucus transport efficiency, whereas regions with lower ciliation showed proportionally reduced clearance capacity. Using these functional ex vivo measurements, we developed quantitative metrics and physics-based computational models capable of predicting mucociliary clearance directly from epithelial composition and ciliary beat activity, providing a framework that links cellular organization to airway activity^25^. The benchmarks and tools established in this study have the potential to enhance the clinical applicability of *in vitro* models for respiratory disease research as they can help assess the effectiveness of treatments for improving/restoring human MCC. For ciliopathies, such treatment will likely rely upon the success of gene-based therapies, including gene replacement, gene editing, and antisense oligonucleotide (ASO) approaches, offer promising avenues to correct the underlying genetic defects in PCD. There has been significant progress in the field in developing gene targeted therapies which offer the potential to restore function at the cellular level^26,27^. However, the success of these approaches depends not only on correcting the genetic defect but also on how many and which cells are corrected. As mucociliary clearance depends on the collective behavior of multiciliated cells at the tissue level, there likely exists a threshold proportion of functional (wild type) ciliated cells required to generate effective, coordinated flow. Below this threshold, even partially corrected epithelia may provide inadequate MCC to significantly impact lung function. Conversely, if only a modest level of correction is sufficient, even partial reconstitution could yield therapeutic benefit.

Beyond informing delivery strategies, knowing the percentage of mutant versus wild-type cells in a chimeric epithelium allows for quantitative modeling of mucociliary dynamics. Such models can predict non-linear, emergent behaviors, where functional gains accelerate beyond a correction threshold, and guide therapeutic benchmarking in preclinical studies. They also provide a framework for correlating molecular and functional biomarkers, such as ciliary beat frequency (CBF), mucus transport rates, or nasal nitric oxide levels, with underlying cellular composition. These insights are critical not only for optimizing therapy, but also for predicting efficacy in clinical trials.

In this study, we focused on CCDC40-mutant PCD cells, a model of severe ciliary immotility, to determine the proportion of wild-type ciliated epithelial cells required to restore effective mucociliary function. Our goal is to provide information toward determining the level of correction required for CCDC40-mutant airways to restore clinically meaningful function in the PCD airway epithelium, thereby supporting the development of scalable and effective molecular therapies that could be built on for all PCD variants.

## MATERIALS AND METHODS

Full details are provided in the Online Supplement

### Experimental Model and Study Participant Details

Deidentified lung tissues for histopathological analysis and HBEC isolation and cryopreserved were obtained from explanted donor lungs either in the Ryan Laboratory with institutional review board (IRB) exemption at the University of Southern California, Los Angeles or obtained with from the University of Iowa Cells and Tissues core in the Precision Medicine Center for Cystic Fibrosis (University of Iowa IRB ID #: 199507432). The study subject had a clinical and genetic diagnosis of primary ciliary dyskinesia due to mutations in CCDC40 and was evaluated a subspecialty clinic directed by Dr. Hornick. HBECs were thawed into tissue culture treated dishes coated with PureCol and expanded in PneumaCult™-Ex Plus complete media.

### Air-Liquid Interface Cultures

CellTracker green CMFDA (1μl, Invitrogen, #C2925) or red CMTPX (1μl, Invitrogen, #C34552) fluorescent dye was added to HBEC cell suspensions. Control HBECs were labelled green and CCDC40 mutant HBECs were labelled red and incubated for 45 mins at 37ºC cells. HBECs were mixed at different ratios of Control: Mutant (100:0, 75:25, 50:50, 25:75, 0:100) and differentiated for a total of 28 days.

### Cilia Beat Frequency (CBF)

CBF was recorded, in the absence of mucus, on a Leica Model DMi8 at 40X, with a region of interest (ROI) of 1024x1024 pixels, for 500 frames with a frame interval of 0.022sec. Cilia beat frequency values are automatically measured by using the open-source ImageJ plugin FreQ.

### Mucociliary Clearance

FluoSpheres™ Polystyrene Microspheres, diluted 1:1000 in complete P-ALI media, were applied directly to the apical side of the insert and immediately images on a Leica Model DMi8 Microscope. One insert per condition per donor was recorded at 10X, with a region of interest (ROI) of 1024x1024 pixels, for 500 frames with a frame interval of 0.022sec. Clearance per beat was calculated using the Track speed (*v* in μm/s) and CBF (*f* in Hz/beats per second) as follows CPB = *v*/*f* (in µm/beat).

### Cilia Length

Differentiated cells were detached from trans-well inserts and applied to a microscope slide with cover slip. 20x images of ciliated cells with distinguishable cell borders and cilia were captured on the Echo Revolution Model RON-K. Cilia length was manually measured in ImageJ with the line tool.

### Quantification and Statistical Analysis

All data were imported into GraphPad Prism for statistical analysis and graph generation. Comparisons between two groups were performed using a *t*-test with Welch’s correction. Comparisons among more than two groups were conducted using one-way analysis of variance (ANOVA) followed by Tukey’s post hoc test to determine statistical significance between groups, unless otherwise specified in the figure legends.

## RESULTS

### Airways with CCDC40 mutations have shorter cilia, increased cellular inflammation and airway occlusion

We obtained airway epithelial cells from a clinically diagnosed PCD patient heterozygous for two pathogenic CCDC40 mutations (c.1620del and c.248del). Prior pathology from this donor reported airway inflammation with variable broncholiths^28^. We further analyzed the specific pathological features associated with large airways (**Fig. 1**). Non-PCD bronchi showed patent airways without inflammation and normal ciliary morphology (**Fig. 1A-B**). CCDC40-mutant bronchi displayed mucus obstruction, airway inflammation, taller epithelial cells, and irregular cilia with neutrophil infiltration (**Fig. 1C-D**). Surface epithelia in PCD tissues had increased height, less regular cilia and neutrophilic exocytosis/infiltration (**Fig. 1D**, arrows). ATUB and RSPH9 staining confirmed comparable multiciliated cell distribution between groups, but cilia were significantly shorter in CCDC40 mutants (**Fig. 1E-F**). Cillia length was measured in cells isolated from wild-type and CCDC40 mutant donor HBECs differentiated at the air-liquid interface (ALI). Wild type multiciliated cells had an average cilia length of 7.68±0.80μm (N=76), that closely matches human cilia length in literature^29^. CCDC40 mutant cells had an average cilia length of 5.42±0.47μm (N= 23), a significant 29% reduction in length compared to the WT counterparts (p<0.0001) (**Fig. 1I-J**). There was no significant difference between the cilia length between independent wild type donors (**Figure E1A**).

**Figure 1:**
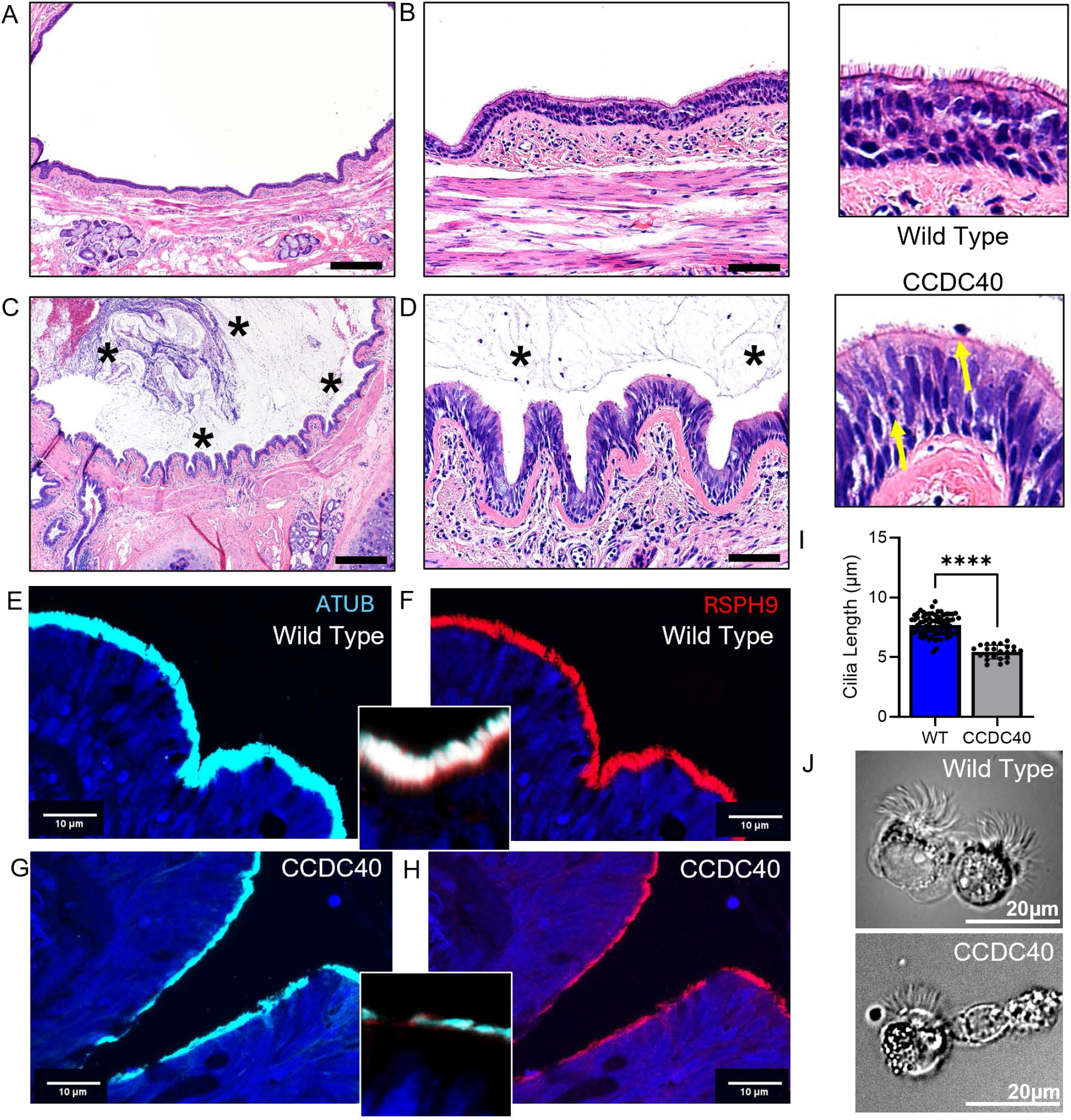
Airway epithelia heterozygous for CCDC40 mutations have severely impacted airway composition and ciliary structure. Representative H&E images showing large airways from a wild type (**A-B**) and a CCDC40 (**C-D**) mutant containing donor with 5x (**B&D**) and 25x (**B&D** right panel images) zoomed images focused on the pseudostratified epithelia. The stars in **C-D** indicate regions variably obstructed with wispy mucus and cellular inflammation (mostly neutrophilic). The yellow arrows in the right zoomed panel **D** highlight inflammatory cell infiltration of the epithelia. Representative immunofluorescent images of the multiciliated cells in the airway epithelium stained for acetylated tubulin (ATUB, cyan) and RSPH9 (red) for wild type (**E-F**) and CCDC40 mutant (**G-H**) tissues. Inset images in **F&H** are 5x zoomed merged images. **I**) Averaged data for ciliary length comparing wild type to CCDC40 mutant cells with representative phase contrast images (**J**). Scale bars in panels **A&C** represent 400 µm, panels **B&D** represent 80 µm. **E-H** represent 10 µm and **J** represent 20 µm. Statistics in panel **I** represent a student’s unpaired t-test N=4 WT and N=1 (n=4) CCDC40 biological donor and 23-26 ciliated cells were evaluated per donor. Supporting data is included in **Figure E1**.

Scanning electron microscopy (SEM) images further highlight the differences in ciliary length between the wild type and CCDC40 donor airways **(Fig. 2A-B**). Wild-type epithelia had dense, uniform cilia, whereas CCDC40 mutants showed reduced density, disorientation, and microvilli-rich regions. **Figure E1B-E** highlights these differences at a variety of SEM magnification levels. WT TEM imaging showed normal 9+2 ultrastructure, while CCDC40 mutants exhibited central pair loss, misoriented or missing outer pairs, and <25% intact 9+2 axonemes (**Fig. 2C-D** and **Figure E1F**). The central pair is absent in over 75% of the basal bodies (**Fig. 2E**) with the majority containing >2 central centrosomes, 6 are present in **Figure 2Di** and 4 in **Figure 2Dii**. In other basal bodies outer pairs are missing and/or disoriented (**Fig. 2Diii**) with <25% of basal bodies having the correct “9+2” assembly (**Fig. 2Div** and **Fig. 2E**). Using PCD Detect to map and overlay the centriole pairs it is clear that the CCDC40 mutant basal bodies are lacking inner dynein arms (**Fig. 2F**). Mutant basal bodies were variably oriented and unevenly spaced, lacking WT planar polarity, as reflected in rose-plot analyses (p<0.001) (**Fig. 2 G-I and Figure E1F**). Mutant cells also displayed profoundly disordered ciliogenesis, with centriole-derived microtubule sets emerging at distant and opposing positions indicative of failed basal body maturation and polarized MT organization (**Figure E1G-I**). Altogether, this data highlights how CCDC40 dysfunction disrupts both the ultrastructural integrity and the coordinated orientation of the multiciliary apparatus, leading to a disorganized ciliary field compared to the wild type multiciliated cells.

**Figure 2:**
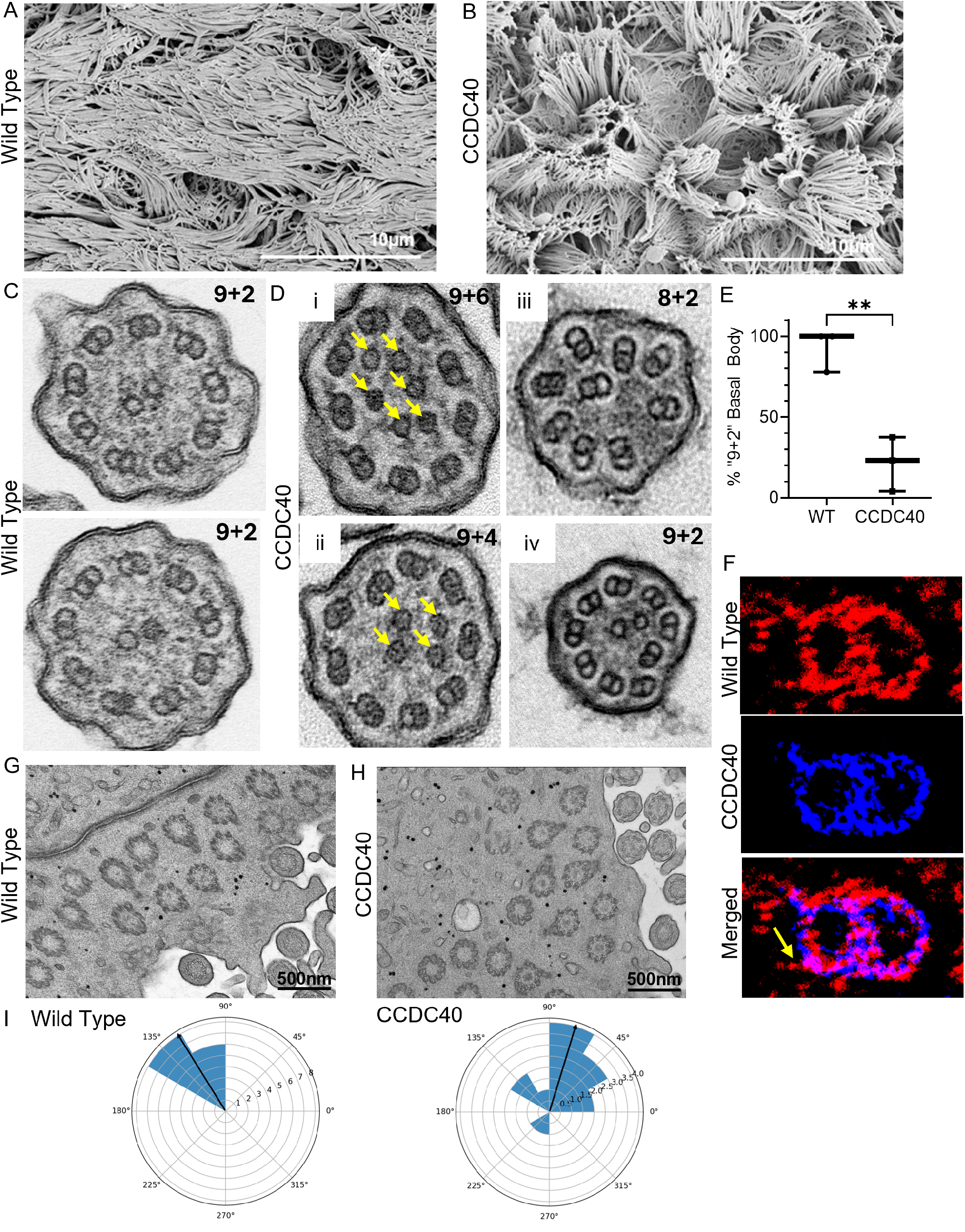
CCDC40 mutations disrupt ultrastructure integrity and coordinated orientation of the multiciliary apparatus. **A-B)** Representative SEM images of cilia in wild type (**A**) and CCDC40 mutant (**B**) differentiated epithelia. **C-D**) Representative TEM images comparing 9+2 centriole arrangements in wild-type multiciliated cells (**C**) to dysregulated arrangements in CCDC40 mutant cilia including 9+6, 9+4 and 8+2 misaligned centriole pairs. Aberrant central centrioles are marked with yellow arrows. **E**) represents the percentage of “normal 9+2” centriole arrangements in the basal bodies comparing wild-type and CCDC40 mutant multiciliated cells. Counts are taken from 3 independent TEM images with ** p<0.01 using unpaired t-test with Welches’ correction. **F**) Overlayed centrioles using PCD Detect, highlighting loss of inner dynein arm in CCDC40-mutants. **G-H**) Representative TEM images highlighting the orientation of the basal feet comparing wild-type (**G**) to CCDC40 mutant cells (**H**). I) Rose plots comparing basal feet directionality in WT and CCDC40 mutants (P<0.01, Watson-Williams test). Scale Bars in A-B represent 10µm, G-H represent 500nm. Supporting data is included in **Figure E1**.

### CCDC40 mutant HBECs maintain full differentiation potential with comparable cellular distribution to wild type HBECs

Airway inflammation and infection may decrease ciliated cell abundance in PCD lungs independently of genotype-determined intrinsic differentiation programs. To directly determine differentiation potential of CCDC40-mutant airway epithelial basal cells in the absence of extrinsic inflammation and infection, CCDC40 donor HBEC were compared to HBEC from 5 independent donor lungs with no prior history of chronic lung disease and no known diagnosis of primary ciliary dyskinesia. The core donor demographics are included in **Table E1**. HBEC from wild-type donors were labelled with cell tracker green and CCDC40 mutant HBEC in cell tracker red to identify the cellular distribution, this was analyzed on Day 0 after the initial ALI seeding of HBEC (**Fig. 3A**) and on the day of air lift (**Fig. 3B**). Proliferation and early distribution did not differ between mutant and WT HBECs. Quantification of the number of red and green cells on the day of air lift represented 1:0.43 (25% red plated), 1:1.14 (50% red plated) and 1:3.23 (75% red plated), not significantly divergent from the original plating ratios (**Fig. 3C**). Additional donor examples are included in **Figure E2**.

**Figure 3:**
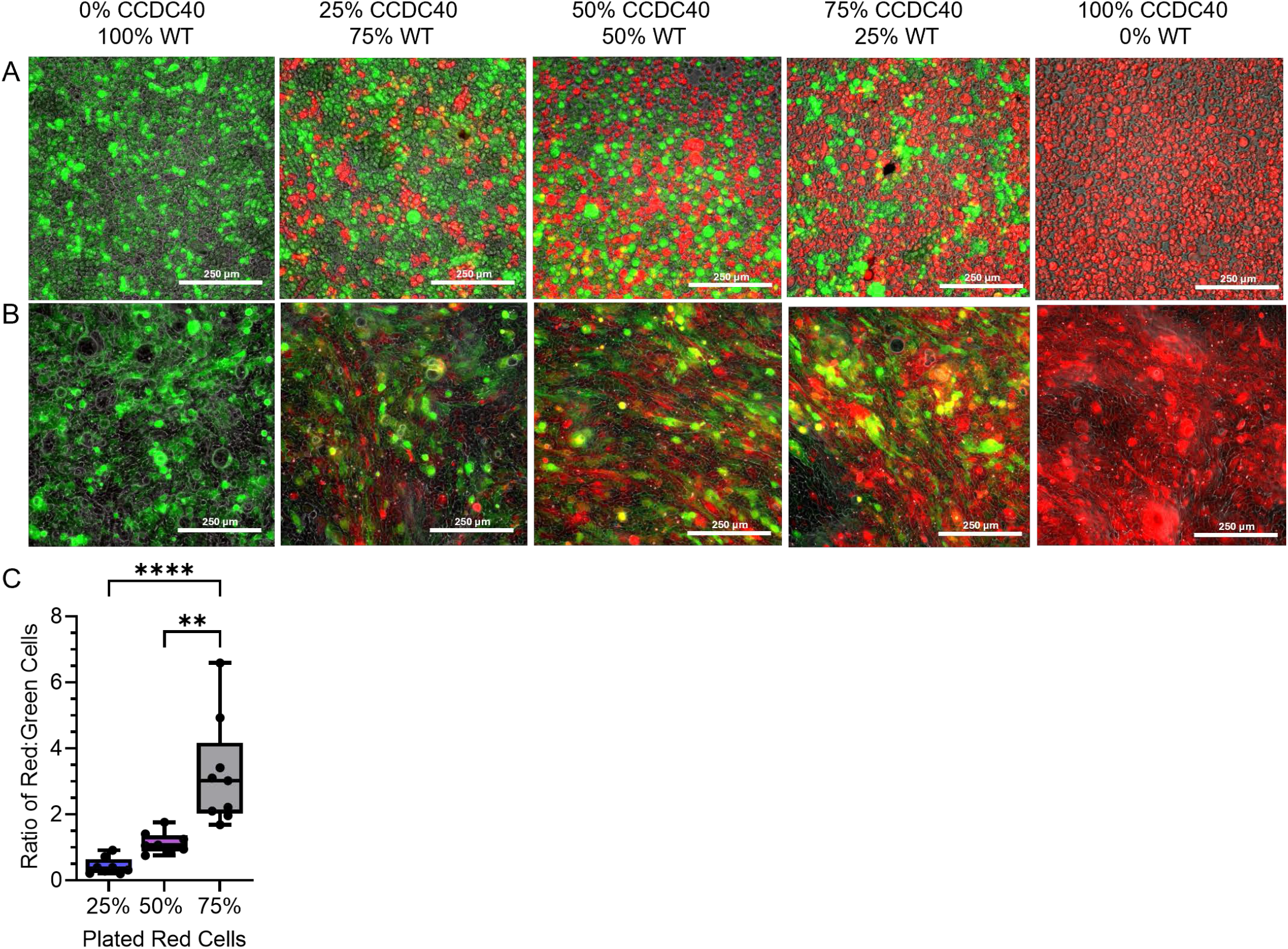
Distribution of different ratios of mutant and wild-type donor HBEC remains similar during pre-ALI expansion. Representative fluorescent images of cells at seeding (Day 0, **A**) and upon airlift (Day 3, **B**). Wild type cells labeled with cell tracker green and CCDC40 mutant cells with cell tracker red. Scale bar represents 250μm in all images. **C)** Averaged ratio of red and green labelled cells at Day 3 (airlift) for all wild-type donors at each plating density, 0% was all red and 100% was all green. Statistics in panel C represent 2way ANOVA with Tukey’s multiple comparisons test **p<0.01 and ****p<0.0001, N=4 Biological donors with two experimental repeats per donor analyzed. Supporting data is included in **Figure E2**.

HBEC were cultured at the ALI to fully differentiated airway epithelia for 28-35 days prior to analysis. Inter donor variability was notable, and data for individual donors is featured in the online data file. Data included in **Figure 4** compares four independent wild-type donors in mixed cultures with our single CCDC40 mutant donor. Efficiency of differentiation to airway epithelium was evaluated by *en face* staining and confocal imaging of the airway culture surface evaluating multiciliated cells (Acetylated alpha tubulin, ATUB), Club cells (Club cell 10kDa protein, CC10) and goblet cells (Mucin 5AC, MUC5AC). Cell boundaries were stained with F-Actin (Phalloidin) (**Fig. 4A**). Cell size trended higher in the CCDC40 donor but varied by donor pairing and was not statistically significant (**Fig. 4B**). However there was a significant decrease in the number of CCDC40 donor cells in each ALI culture, supporting the trend to increased cell size (**Fig. 4C**). Ciliated coverage decreased in the 100% mutant cultures, with corresponding increases in goblet cell coverage (**Fig. 4D**). Accordingly, the surface coverage of Goblet cells increased with significant differences in the 75 and 50% (p< 0.05)compared to the 100% CCDC40 mutant (**Fig. 4E**). CCDC40 RNA was significantly reduced in the CCDC40 mutant donor cultures (**Fig. 4F**). While there was significant donor heterogeneity in gene expression across the wild type donors there was also a significant increase in early multiciliogenesis gene MCIDAS and inflammatory gene TNFα (**Fig. 4F**). Other genes tended to decrease but were not statistically significant, including CCNO and ODF2 (data not shown).

**Figure 4:**
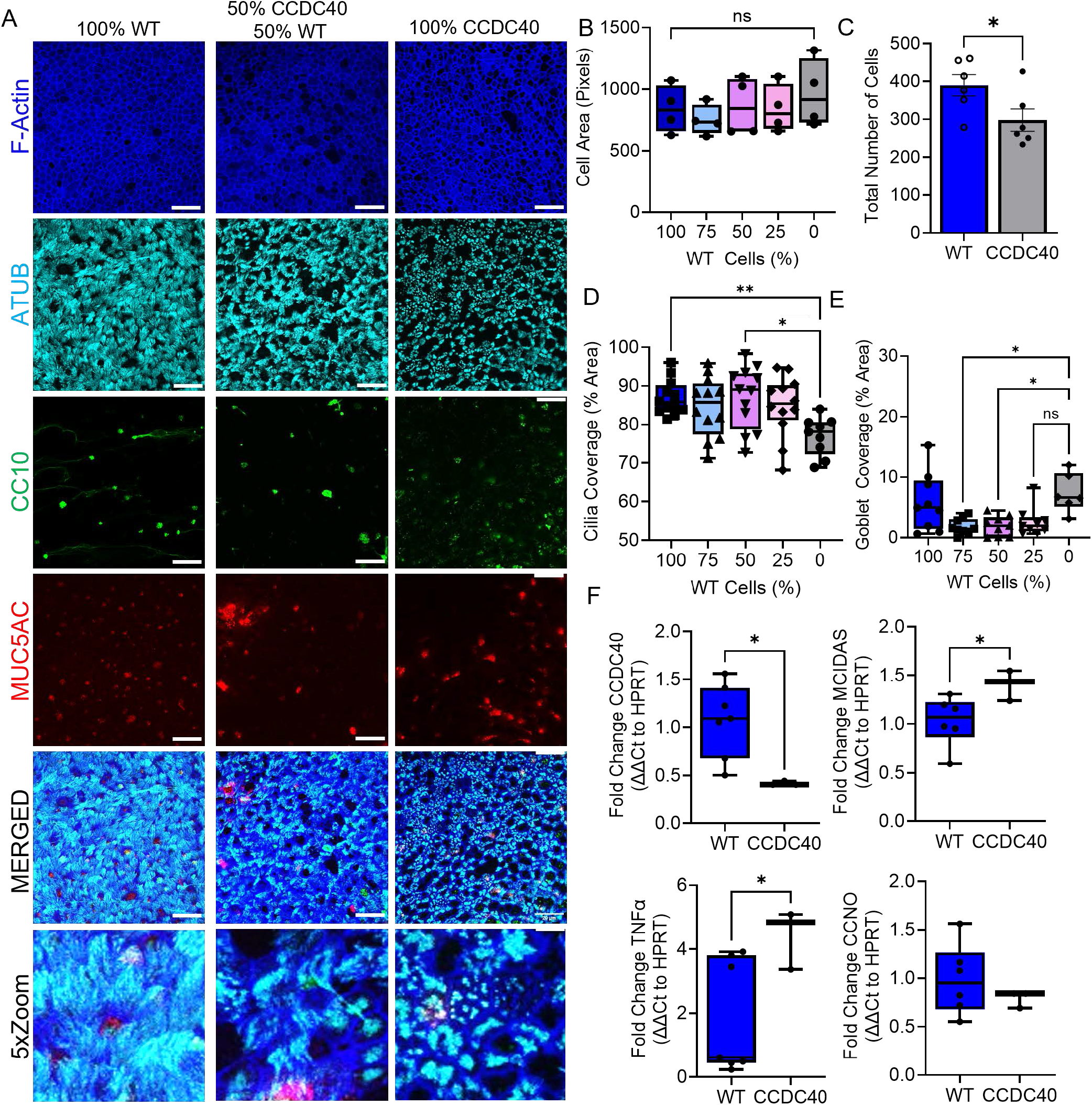
Patterning of differentiation in CCDC40 mutant cells favors larger cells with a lower ciliated cell density. **A)** Representative immunofluorescent images of the differentiated HBEC cultures from Donor 4 and the CCDC40 mutant donor cells, each at 50% and 100% of the culture, stained for acetylated tubulin (ATUB, cyan), club cell 10kDa protein (CC10, green), mucin 5AC (MUC5AC, red) and phalloidin (F-ACTIN, blue). Scale bars in all panels represent 20 µm in all images and the lower images are 5x zoomed merged images. Averaged cell area per cell (**B**), total number of cells comparing WT to CCDC40 (**C**), ciliated cell surface area (**D**) and secretory cell surface area (**E**) comparing all ratios of wild type to CCDC40 mutant cells. **E**) qRT-PCR comparing WT (4 biological donors) to CCDC40 mutant donor (4 experimental repeats). Statistics in panel **B, D&E** represent a 2-way ANOVA with Tukey’s multiple comparisons test **p<0.01 and ****p<0.0001, N=4 Biological donors with two experimental repeats per donor analyzed. Statistics in **C&F** panels represent an unpaired t-test with Welch’s correction, *p<0.05, N=4 Biological donors with two experimental repeats per donor analyzed. Supporting data is included in **Figure E3**.

### Functional mucociliary clearance and ciliary dynamics are impaired by increasing presence of CCDC40 mutant HBEC in mixed cultures

In addition to cellular composition, functional mucociliary clearance is determined by ciliated cell abundance and activity. Moreover, cells with impaired ciliary function may affect neighboring cells via contact-dependence or paracrine signaling abnormalities. We therefore hypothesized that mucociliary clearance is impacted by both autocrine and paracrine signals decreasing in a non-linear trend with increased presence of mutant cells. To evaluate this we quantified ciliary function and mucociliary clearance, determining the impact CCDC40 mutant cell abundance had on tissue function **(Fig. 5**). Ciliary activity became increasingly patchy with higher mutant percentages, and bead tracking demonstrated progressive loss of coordinated flow (**Fig. 5A-B and Figure E3A**). The mean particle clearance speeds across all donors were significantly decreased with increasing presence of donor cells from ∼56.5 μm/s at 100% WT to ∼26.5 μm/s in 25% WT (**Fig. 5C and Figure E3B**), with track speeds in the mutant-donors cells moving minimally at ∼9.4 μm/s.

**Figure 5:**
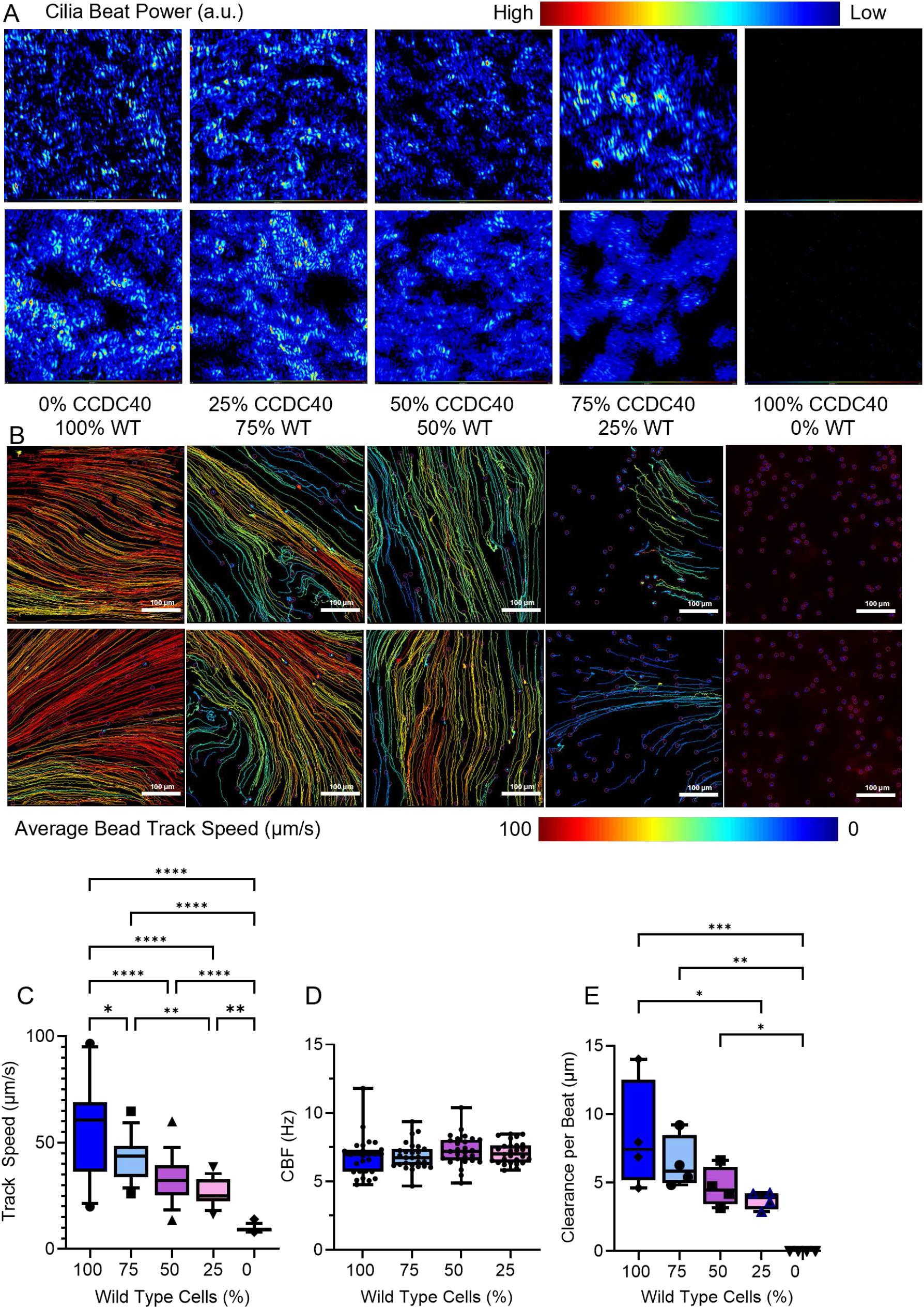
Distribution of WT and CCDC40 cells regulates functional trajectories. **A**) Cilia beat power heat map overlay from CBF recordings displayed in arbitrary units. Heat map legend provided above. Larger power output is designated by warmer colors; lower power output designated by cooler colors. **B**) Representative images of tracked fluorescent bead flow on human airway epithelium. Tracks are color coded based on particle speed in µm/sec. 100µm scale bar included. **C**) Box and whisker plot of measured particle clearance speed from tracked fluorescent bead movement. **D**) Box and whisker plot of average cilia beat frequency in hertz **E**) Box and whisker plot of calculated ciliated cell particle clearance per beat in µm. Scale bars in all panels represent 100 µm. *p < 0.05, **p < 0.01 by one-way ANOVA with post-hoc Tukey’s test (**C** and **E**). n=8 technical replicates from 5 biological replicates for each experiment. Supporting data is included in **Figure E3**.

Across donors, WT ciliary beat frequency remained unchanged despite increasing mutant representation **(Fig. 5D**). CBF measurements were not included for CCDC40 mutant controls due to the absence of motile cilia in these cultures. Analysis of individual donor responses revealed variable CBF changes with increasing proportions of mutant cells **(Figure E3C**). While some donor lines exhibited marked increases in frequency, others showed little change; however, none demonstrated a decrease in CBF. To further assess ciliary function, we calculated the clearance per beat (CPB) as a measure of clearance efficiency (**Fig. 5E**). CPB increased significantly with the proportion of ciliated WT cells, starting from 0 µm/beat (undetectable) in 0% WT (100% mutant) populations. Cultures containing 25–50% WT cells exhibited average CPB values of 3.7–4.6 µm/beat, while those with 75–100% WT cells reached 6.4–8.4 µm/beat.

Consistent with the reductions observed CPB, the directional coordination of flow also declined with increasing proportions of mutant cells. Autocorrelation, a measure of the directional persistence of particle motion ranging from 0 (random movement) to 1 (fully synchronized motion), reflected this trend. Autocorrelation was highest in 100% WT cultures and declined progressively with more mutant cells (**Fig. 6A**). Cultures containing 75–50% WT cells showed modestly lower averages of ∼0.40, while 25% WT populations exhibited a sharper and significant decrease to ∼0.14 and a narrower spread in data variability. In 0% WT cultures, autocorrelation approached zero (average ∼0.04), indicating a near-complete loss of coordinated flow. Again, there is significant individual donor variability in the autocorrelation values at 100% WT, however the data consistently decreased with increasing mutant cell proportions, becoming more uniform and compressed across donors. Representative magnitude and vector XY plots of bead trajectories further illustrate the progressive loss of clearance directionality and coordination across WT-to-mutant ratios (**Fig. 6B-C**).

**Figure 6:**
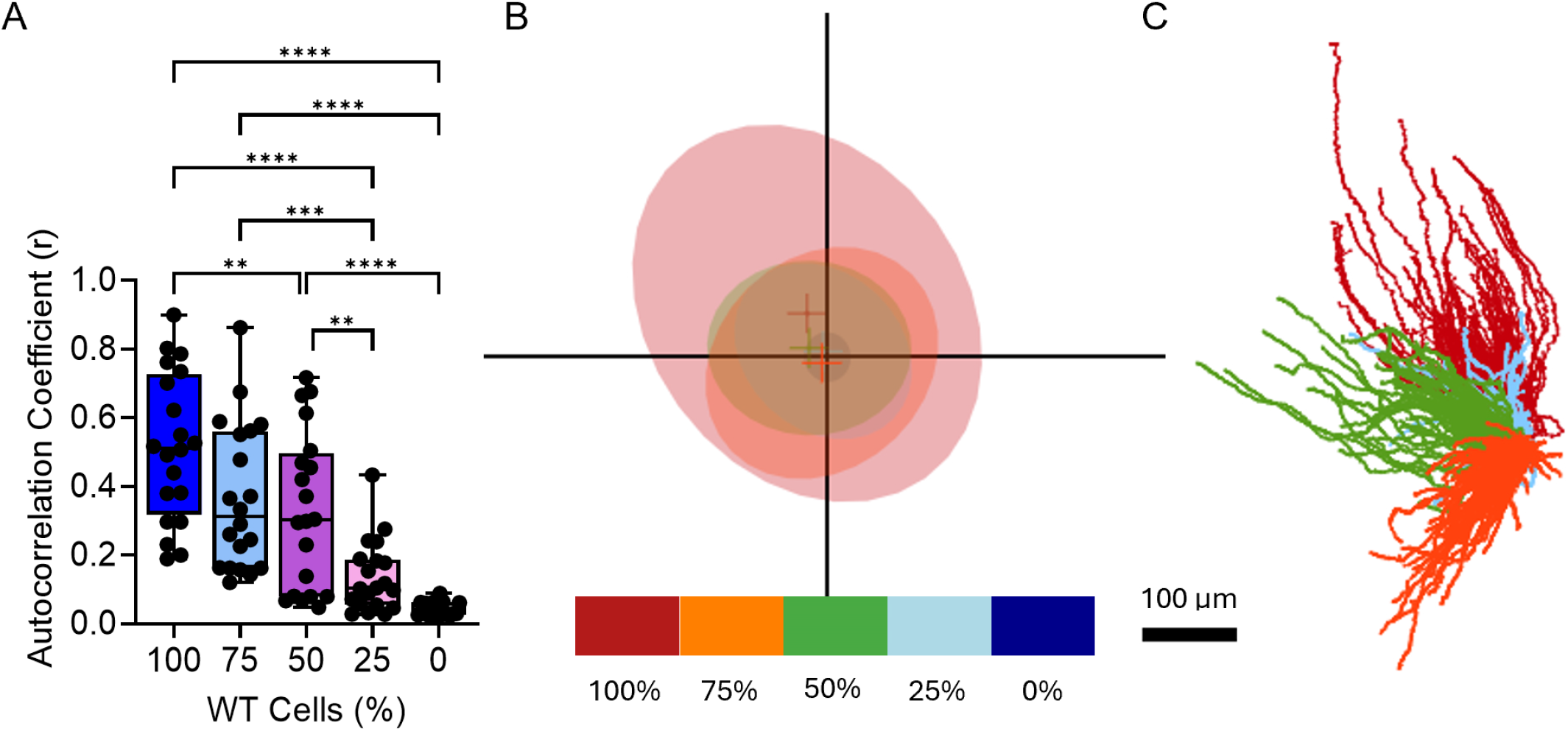
Characterizing directionality and persistence. **A**) Compiled box and whisker plot of average autocorrelation values of tracks within the first 6 seconds of fluorescent bead flow. **p <0.01, ***p < 0.001, ****p < 0.001 by one-way ANOVA with post-hoc Tukey’s test (**C** and **E**). n = 20 regions per condition from 4 biological replicates for each experiment. **B**) Representative plot of magnitude and directional persistence of fluorescent bead flow. Each ellipse, color coordinated by percent wild type as depicted in legend below, represents the average speed and directional preference of tracks associated with one FOV of fluorescent bead flow. Ellipses are plotted on a directional plane to visualize movement persistence in a particular direction from the origin of bead flow. **C**) Bead flow tracks from donor 1 associated with subfigure A, color coded the same.

To predict the impact of changes in ciliary coverage and mucociliary clearance, we examined how CPB scaled with the proportion of ciliated WT cells. In our prior work we have established structure–function principles investigating coordinated ciliary organization and sufficient coverage as critical determinants of effective mucociliary transport to predict in vivo clearance from in vitro tissue models^30^. Using tissue-level metrics shown to predict mucociliary clearance in vivo, we plotted CPB values against the total number of wild-type ciliated cells in each culture (**Fig. 7A**) and against the total number of ciliated cells regardless of genotype (**Fig. 7B**). Interestingly, when CPB was plotted as a function of the percentage of WT-only ciliated cells, CPB consistently performed at or above the expected efficiency threshold **(Fig. 7A)**. This suggests that the proportion of functional WT ciliated cells, rather than the total ciliated cell burden, is the dominant driver of restored mucociliary performance. When considering the total percentage of ciliated cells (mutant + wild type), our data indicate that cultures containing >75% WT cells within the ciliated population aligned with human ex vivo mucociliary clearance metrics (**Fig. 7B**, square symbols). In one donor, even 50% WT cells (Blue open triangle, Donor 1) met these performance benchmarks, with 75% and 100% WT conditions exceeding predicted efficiencies. A one-phase association model (R^2^ = 0.71) revealed a performance plateau at approximately 7 µm/beat, reached once cultures contained ∼60–70% WT ciliated cells. Although variability between donors increased at higher WT proportions, the overall relationship remained robust. The curve midpoint occurred at ∼32% WT ciliated cells, with an estimated CPB of ∼5.65 µm/beat at 50% WT cells **(Fig. 7C)**. Together, these findings reveal a nonlinear restoration profile, in which mucociliary performance improves once 30–40% of the ciliated cells are wild-type and begins to plateau as WT representation approaches two-thirds of the ciliated population. This nonlinear behavior underscores a threshold requirement for functional rescue and highlights the disproportionate contribution of WT cells to coordinated clearance.

**Figure 7:**
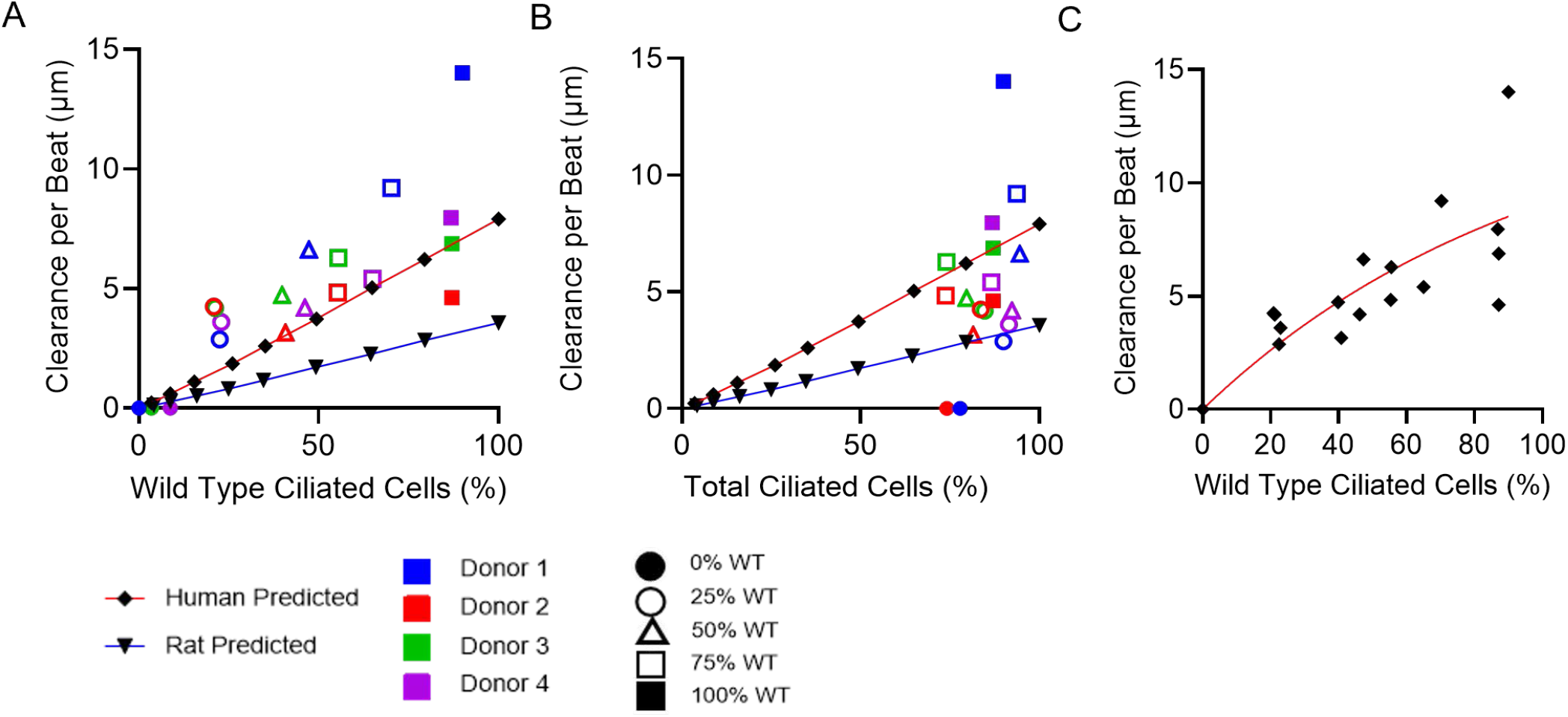
Non-linear threshold levels of functional cilia required for effective transport modelling. **A**) CPB plotted against percent WT ciliated cells per donor & condition, as well as predicted models for Human and Rat airway CPB (published in ^30^) against ciliated cell percentage. Colors denote donor line; shapes denote percent WT input condition. Rat airway prediction (Blue line), Human airway prediction (Red line). **B**) Same graph as figure **6A**, with donor raw ciliated cell percentage instead of calculated WT ciliated cell percentage. **C**) Fit curve analysis of CPB using a one-phase association model (R^2^ = 0.71). Each point represents average clearance per beat from one donor, derived from eight replicate videos. Fit parameters were estimated without constraints.

## DISCUSSION

In this study, we established a quantitative framework to define how the proportion of functional epithelial cells impacts mucociliary clearance in CCDC40-associated PCD. Using primary HBEC cultures, we modeled chimeric epithelia containing WT/CCDC40-mutant ratios, enabling precise evaluation of ciliary structure and function across genotypic gradients. Three key insights emerged: 1) CCDC40 mutations shorten cilia, disrupt coordinated beating, stall clearance, and increase epithelial inflammation; 2) mucociliary transport is highly sensitive to losses in the functional ciliated-cell fraction, with directional clearance collapsing below a threshold; and 3)∼50–75% WT cells among ciliated cells restores near-normal clearance, providing a quantitative benchmark for therapeutic correction in CCDC40-mutant airways.

Consistent with prior reports, CCDC40 deficiency produced ultrastructural and functional defects including, short, immotile cilia and dynein arm disorganization^22,24^. The ∼25% reduction in cilia length aligns with prior analyses of CCDC39/40 loss affecting axonemal architecture and microtubule doublet orientation^22^. Because mucociliary clearance relies on synchronized MCC that generates directional flow^25^ transport declined nonlinearly as mutant representation increased: reduced WT coverage yielded coordinated decreases in CPB and flow alignment, underscoring the cooperative, emergent nature of cilia-driven transport. Modeled against the total number of ciliated cells, ≥∼75% WT within the ciliated population was required to approach ex vivo human airway efficiency, an effect that must be interpreted within a PCD-dominated epithelium where dyskinetic/immotile cilia act as mechanical sinks that add drag, distort local flow fields, and impede WT cells’ ability to entrain collective transport. This distinction is evident when contrasted with settings lacking disruptive mutant neighbors: several in vitro and in vivo observations indicate that lower fractions of functional ciliated cells (∼30–60%) can sustain normal or only mildly reduced clearance when the remaining cells do not introduce strong counter-flows or drag (e.g., heterozygous DNAAF6 carriers with mosaic functional ciliation)^31^. Likewise, disease-modeling studies show that partial restoration of ciliary coordination improves flow even without uniform correction, highlighting that mucociliary systems possess redundancy when non-functional cells are not actively disruptive^32^. Together, these comparisons reinforce that the elevated threshold observed here is not due to a necessity for high ciliary coverage per se but rather reflects the deleterious biomechanical contribution of PCD cells.

These relationships carry direct therapeutic implications. Partial gene correction may not translate linearly into functional rescue; clinically meaningful benefit may require restoring function in a critical mass of ciliated cells, estimated here at ∼75% of the ciliated population (**Fig. 6B**). Because only a subset of epithelial cells differentiate into ciliated cells at steady state, achieving that level of correction among ciliated cells likely demands efficient transduction of a large fraction of the epithelial population. Vector systems with strong tropism for basal progenitors could support durable correction across epithelial turnover, while airway-scale flow modeling can anticipate how heterogeneous correction shapes regional clearance and guide dosing and vector distribution.

Genotype specificity is also expected. Different PCD mutations (impacting axonemal structure, beat pattern, or coordination) probably impose distinct “negative MCC values,” shifting the correction thresholds for functional rescue. Systematic assessment across PCD genotypes could map mutation-specific baseline thresholds and generate a sharable mutation-phenotype-threshold atlas. More broadly, our findings provide measurement frameworks, high-resolution structural assessment, single-cell genotype/function coupling, and quantitative MCC, that are already feasible on bronchoscopic or nasal mucosal biopsies, supporting dose selection and pharmacodynamic evaluation aligned with regulatory expectations.

Together, these data delineate the structural and functional signatures of CCDC40-mutant airway epithelia and quantify a PCD-specific correction threshold for restoring mucociliary clearance to ex vivo human tissue benchmarks. Building on our prior work they also offer experimental and computational tools for benchmarking gene-therapy efficacy and guiding clinical translation. Future work should extend this analysis across PCD genotypes to refine mutation-specific thresholds, incorporate in vivo validation with engineered airway models, and examine the temporal dynamics of epithelial repair after partial correction. Ultimately, clarifying how cellular composition governs coordinated ciliary motion, while accounting for the disruptive mechanics of mutant cilia, will be essential to advancing curative PCD therapies.

## ACKNOWLEDGEMENTS

We would like to thank you to Greg Bonde for his mentorship of the undergraduate students who did the research in this study.

## RESOURCE AVAILABILITY

### Lead contact

Further information and requests for resources and reagents should be directed to and will be fulfilled by the lead contact, Amy Ryan.

### Materials availability

Due to limited expansion of primary HBEC the donor derived CCDC40 mutant cells are not able to be shared. Cells from donors with no prior history of lung disease are available from the corresponding author and through the Cells, Tissues and Models Core within the Precision Medicine Center for Cystic Fibrosis (CF).

### Data and code availability

All raw data including qPCR data files and excel files with CBF and MCC metrics for all donors and repeats and Python code for creating rose plots is deposited at Figshare DOI: 10.6084/m9.figshare.30407797 and are publicly available as of the date of publication. Microscopy data reported in this paper will be shared by the lead contact upon request.

## SUPPLEMENTAL INFORMATION

The Supplemental Document contains main Supplemental Materials and Methods and Figures E1–S3, Tables E1-E3 and supporting Movie Files E1-S4.

**Table E1:** Human bronchial epithelial cell (HBEC) information

**Table E2:** Primer pairs used for gene expression analysis

**Table E3:** Antibodies used for IF Staining

**Figure E1:** Supporting Figure 1 & 2: **CCDC40 mutant donor-derived ciliated cell have shorter, less uniform cilia. A**) Ciliary length per donor, each for represents an individual field of view from a minimum of three independent ALI differentiations. Data represents mean ± SEM, one way ANOVA ****p<0.0001. **B-E**) SEM images for CCDC40 mutant and Wild Type donors shown at two different magnifications. **F**) TEM images highlighting abnormal centriole replication and arrangements and misalignment of ciliary basal feet. **G**) TEM images highlighting unusual centriole-derived microtubule sets emerging at distant and opposing positions in enlarged structures.

**Figure E2:** Supporting Figure 3: **Distribution of different ratios of mutant and wild-type donor HBEC remains similar during pre-ALI expansion**. Representative IF images of Cell Tracker Red labelled CCDC40 mutant HBEC and CellTracker Green labelled WT donor HBEC for an additional two donors. **A** and **C** are shown at day 0 of airlift and **B** and **D** are shown after 3 days of airlift across all plating ratios. Scale bars in all images are 250µm.

**Figure E3:** Supporting Figure 5: **Distribution of WT and CCDC40 cells regulates functional trajectories**. Normalized track speed (**A**) measured particle clearance speed from tracked fluorescent bead movement (**B**), CBF (**C**) and autocorrelation co-efficient (**D**) with lines representing each donor. Data represents average of technical replicates from 5-6 independent biological replicates for each experiment.

---

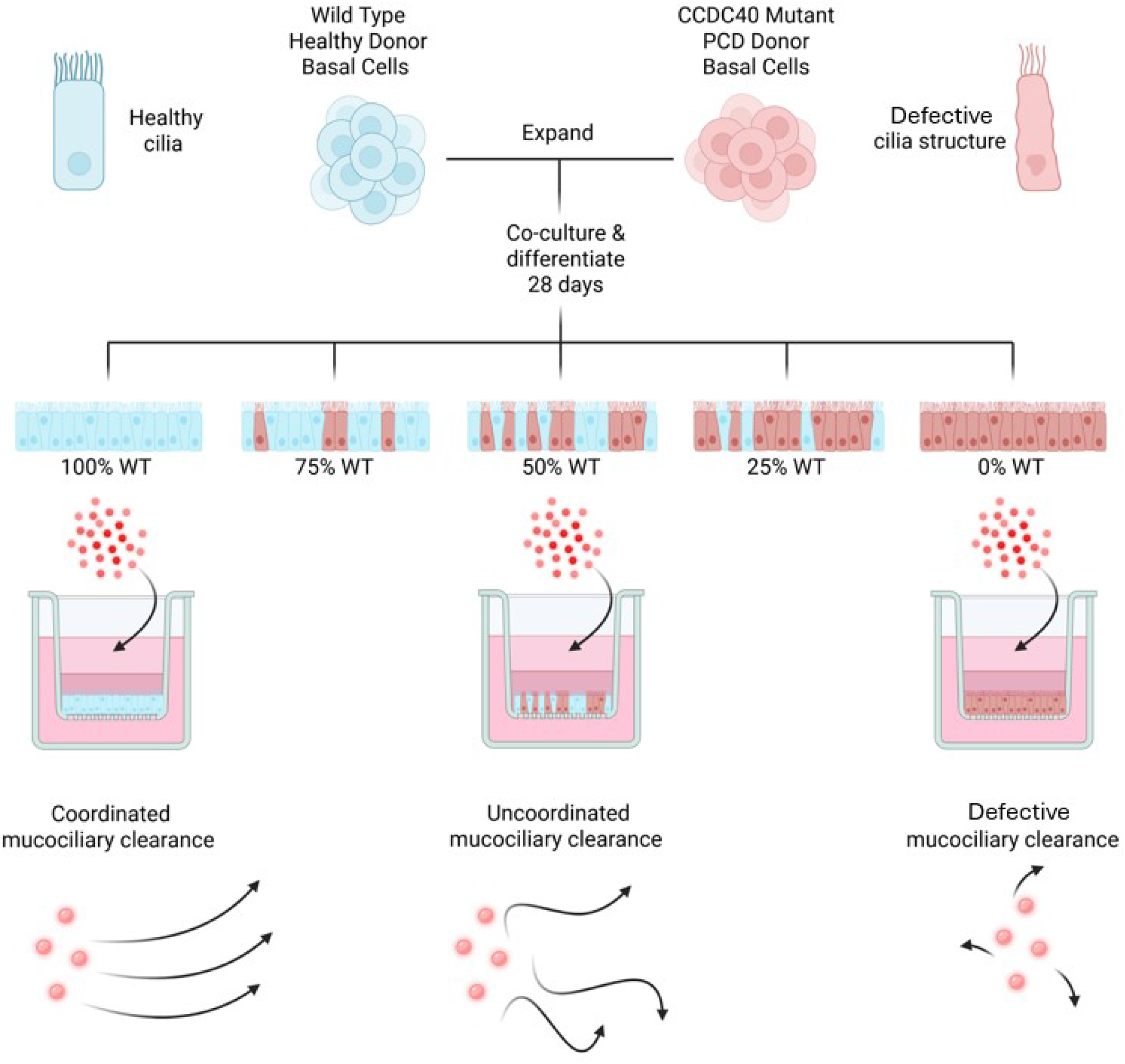

