## Supplemental Data File for "Defining Functional Correction Thresholds in Primary Ciliary Dyskinesia for Effective Gene Therapies"

**Author Contributions:** Conceptualization, A.L.R. and E.J.Q.; methodology, A.L.R., B.E.F., B.J.W., J.E.B., E.J.Q., E.C.L., L.K.G., T.M. and D.K.M.; Investigation, A.L.R., B.E.F., B.J.W., J.E.B., E.J.Q., E.C.L., L.K.G., A.A.P., T.M. and D.K.M.; writing—original draft, A.L.R., B.E.F., B.J.W., and J.E.B., writing—review & editing, A.L.R., B.E.F., B.J.W., J.E.B., E.J.Q., E.C.L., L.K.G., A.A.P., T.M., D.B.H. and D.K.M.; funding acquisition, A.L.R.; resources, A.L.R.; supervision, A.L.R.

**Sources of Support:** This study was supported by the National Institutes of Health (NIH):NHLBI R01 HL153622 (ALR). The cells were provided through the Cells, Tissues and Models Core supported in part by the NIDDK P30 DK054759 (ALR).

**Running Head**: Impact of CCDC40 mutations of Mucociliary Clearance

### SUPPLEMENTAL MATERIALS AND METHODS

**Experimental Model and Study Participant Details**: Deidentified lung tissues for histopathological analysis and HBEC isolation and cryopreserved were obtained from explanted donor lungs either in the Ryan Laboratory with institutional review board (IRB) exemption at the University of Southern California, Los Angeles or obtained with from the University of Iowa Cells and Tissues core in the Precision Medicine Center for Cystic Fibrosis (University of Iowa IRB ID #: 199507432). The study subject had a clinical and genetic diagnosis of primary ciliary dyskinesia due to mutations in CCDC40 and was evaluated a subspecialty clinic directed by Dr. Hornick. HBECs were thawed into tissue culture treated dishes coated with PureCol (Biomatrix, 5005-100ML) diluted 1:30 and expanded in PneumaCult™-Ex Plus complete media (PEx+, StemCell Technologies, #05041) with 0.2% primocin (InvivoGen, #ant-pm-2) and 0.1% hydrocortisone (Sigma, #H0135-1MG). Media was replaced every 48 hours. At ~80% confluency, cells were split by resuspension using Accutase (Innovative Cell Tech, #AT 104-500), collected by centrifugation at 400x g, 5 min, and resuspended in 1mL of DMEM/F12 (Gibco, #11320-033), counted and plated at $5x10^3$ cells/$cm^2$.

**Air-Liquid Interface Cultures**: Celltracker green CMFDA (1µl, Invitrogen, #C2925) or red CMTPX (1µl, Invitrogen, #C34552) fluorescent dye was added to HBEC cell suspensions. Control HBECs were labelled green and CCDC40 mutant HBECs were labelled red and incubated for 45 mins at 37ºC cells. Labelled cells were washed by centrifugation at 400x g, 5 min, resuspended in 1ml of PEx+ and counted using a hemocytometer. Cells were then prepared at 750,000 cells/ml and using 5 separate 1.5ml Eppendorf tubes, control and mutant HBECs were mixed at different ratios of Control: Mutant (100:0, 75:25, 50:50, 25:75, 0:100). Cell co-cultures were seeded at 75,000 cells per 6.5mm trans-well insert (Corning, #3470), prepared with PureCol coating. Cells seeded on trans-well inserts were expanded in PEX+ media (500ul basolateral, 100ul apical) for 3 days, changing media on the 1$^{st}$ and 3$^{rd}$ day. After 3 days, switched to PneumaCult™- Air Liquid Interface Basal Medium (StemCell Technologies, 05002) (P-ALI) + PneumaCult™-ALI 10X Supplement (StemCell Technologies, 05003), PneumaCult™-ALI Maintenance Supplement, 0.4% primocin (InvivoGen, ant-pm-2), 0.2% heparin (Sigma, H3149-100KU), 0.5% hydrocortisone (Sigma, H0135-1MG). 500ul of media added to basolateral side only. Immunofluorescent images of the cells were taken using the Echo Revolution Model RON-K on day 0 immediately after seeding, and on day 3 when the cells were air-lifted to document the distribution of cells. Media was replaced every 48 hours and cultures differentiated for a total of 28 days.

**Cilia Beat Frequency:** To record cilia beat metrics in the absence of mucus, inserts were washed twice with DPBS (Gibco, 14190-144) for 10 minutes at 37C. Recorded inserts are moved to a separate plate, one at a time, to mitigate time spent outside of incubator conditions. One insert per condition were then recorded on a Leica Model DMi8 at 40X, with a region of interest (ROI) of 1024x1024 pixels, for 500 frames with a frame interval of 0.022sec. 8 FOV's were captured per insert (only 6 FOV's captured for Donor 1 inserts). Cilia beat frequency values are automatically measured by using the open-source ImageJ plugin FreQ[1,2]. The average frequency (Hz) value was used for cilia beat frequency figures and clearance per beat values.

**Mucociliary Clearance:** To record ciliary clearance of particles in the absence of mucus, inserts are washed twice with DPBS (Gibco, 14190-144) for 10 minutes at 37C. Recorded inserts are moved to a separate plate, one at a time, to mitigate time spent outside of incubator conditions. FluoSpheres™ Polystyrene Microspheres (Invitrogen, F13083), diluted 1:1000 in complete P-ALI media, are applied directly to the apical side of the insert and immediately taken to a Leica Model DMi8 Microscope. One insert per condition per donor was recorded at 10X, with a region of interest (ROI) of 1024x1024 pixels, for 500 frames with a frame interval of 0.022sec. 8 FOV's were captured per insert (only 6 FOV's captured for Donor 1 inserts). Bead flow tracks (TrackMate): Fluorescent bead flow videos were exported to the TrackMate plugin on ImageJ[2,3]. Fluorescent beads were tracked and filtered for light intensity, and track lengths over 40 microns. Tracks were then colored for mean track speed from a range of 0 to 100 microns per second. Representative tracked images were exported for all FOV's. Motility Lab spreadsheets were exported for further bead flow quantitative analysis. Motility lab is a free browser program that comparatively and quantitatively analyzes exported spreadsheets from TrackMate[4]. From the bead flow FOV's, data for mean track speed, autocorrelation, and directionality plots of tracks were generated. Clearance per beat was calculated using the Track speed ($v$ in µm/s) and CBF ($f$ in Hz/beats per second) as follows CPB = $v/f$ (in µm/beat).

**Cilia Length**: Differentiated cells were detached from trans-well inserts by applying pre-warmed Accutase (Innovative Cell Tech, AT 104-500) apically for 20 minutes, occasionally tapping the plate and pipetting up and down. The Accutase/cell mixture was diluted 1:10 with DPBS (Gibco, 14190-144) and applied to a microscope slide with cover slip. On an Echo Revolution Model RON-K, 10x images of ciliated cells with distinguishable cell borders and cilia were captured. Cilia length was manually measured in ImageJ with the line tool. Representative 40x images were also captured.

**RNA Isolation and qRT-PCR**: Total RNA from two donors (1,2) was isolated using the Direct-zol RNA miniprep Kit (Zymo Research) manufacturer protocol. 300ul of TRIzol was used to lyse cells from ALI insert and was processed with 400ul of Direct-zol RNA PreWash, 700ul of RNA Wash Buffer and eluted into a final volume of 20ul of DNase/RNase free water. All A260/280 values fell between 2-1.7. Total RNA from three donors (3,4,5) was isolated using the TRIzol Reagent (Invitrogen) manufacturer protocol. 500ul of TRIzol was used to lyse cells from ALI insert and was processed with, 100ul of chloroform, 250ul of Isopropyl Alcohol, 500ul of 75% Ethanol and reconstituted in 30ul of DNase/RNase free water. For cDNA synthesis, high-capacity cDNA reverse transcriptase (RT) was performed using Multiscribe Reverse Transcriptase (Thermo Fisher Scientific) according to manufacturer's protocol. 30 µl reactions were created using measured amounts of 10x RT buffer, 25x dNTP mix, 10x Random Primers, DNase/RNase free water and ran through thermocycler at protocols conditions. Quantitative PCR was performed using the Power Up SYBR Green master mix, following manufactures protocol (Thermo Fisher Scientific). PCR conditions were 50C for 2 minutes then 95C for 10 minutes, followed by 40 cycles of 95C for15s and 60C for 60s. Statistical analysis of the PCR was done in Prism using a one-way ANOVA test matching means of each condition with each other. Primers are included in **Table E2.**

**Immunofluorescence staining:** Cells were fixed with 4% paraformaldehyde (PFA) for 15 minutes at room temperature and washed with PBS. Cells were permeabilized using 0.05% Triton-X100 for 5 minutes and then blocked in PBC containing 5% normal donkey serum and 2 % BSA for 1-2 hours. Cells were incubated in primary antibodies (listen in **Table E3**) for 2 hours at room temperature, washed 3 x 5 minutes in PBS and subsequently incubated in secondary antibodies for 1 hour at room temperature. Cells were mounted using Fluomount G (Southern Biotech) and imaged on the Revolution microscope (Echo, A Bioc company). Images were collected with a Zeis LSM 980 confocal microscope. Maximum intensity projections were processed using ImageJ software and images represented as single or merged.

**Scanning Electron Microscopy (SEM):** Samples designated for scanning electron microscopy (SEM) were fixed overnight in 2.5% glutaraldehyde prepared in 0.1 M sodium cacodylate buffer. The samples were rinsed three times in 0.1 M sodium cacodylate buffer for 10 min per rinse, followed by post-fixation in 1% osmium tetroxide for 1 h. Samples were then rinsed again three times in 0.1 M sodium cacodylate buffer for 10 min each and briefly rinsed in distilled deionized water for 1 min. Dehydration was performed using a graded ethanol series consisting of 25%, 50%, and 75% ethanol in water for 15 min each, followed by 95% ethanol and two changes of 100% ethanol for 20 min each. Samples were subsequently treated twice with hexamethyldisilane for 10 min per treatment and allowed to air-dry overnight at room temperature in an incubator. Dried samples were mounted onto SEM stubs using carbon adhesive tabs. Silver paint was applied around the sample edges to enhance conductivity and allowed to dry. Samples were sputter-coated using a K550 Emitech sputter coater and imaged with a Hitachi S-4800 scanning electron microscope.

**Transmission Electron Microscopy (TEM):** Samples designated for TEM were fixed overnight in 2.5% glutaraldehyde prepared in 0.1 M sodium cacodylate buffer. Samples were rinsed three times in 0.1 M sodium cacodylate buffer for 10 min per rinse and post-fixed in a solution containing 1% osmium tetroxide and 1.5% potassium ferrocyanide for 2 h. Following post-fixation, samples were rinsed three times in 0.1 M sodium cacodylate buffer for 10 min each and briefly rinsed in distilled deionized water for 1 min. Samples were stained *en bloc* with 2.5% uranyl acetate for 15 min, followed by dehydration through a graded ethanol series consisting of 50% and 75% ethanol for 15 min each, 95% ethanol for 20 min, and two changes of 100% ethanol for 20 min each. Infiltration was performed using a graded ethanol–epoxy resin series (Eponate 12), consisting of a 2:1 ethanol-to-epoxy mixture for 30 min, a 1:2 ethanol-to-epoxy mixture for 1 h, and two changes of 100% epoxy resin for 30 min each. Samples were then embedded in fresh epoxy resin and polymerized overnight at 60°C. Ultrathin sections (~70–90 nm) were cut using an ultramicrotome (Leica EM UC6) equipped with a DiATOME diamond knife and collected onto copper TEM grids. Sections were post-stained with uranyl acetate and lead citrate prior to imaging. Imaging was performed using a transmission electron microscope (Hitachi HT7800), operated at an accelerating voltage appropriate for biological specimens (80kV).

**EM Analysis**: TEM cross-sections of cilia were processed using PCD Detect[5], a software toolkit that enhances ultrastructural features through image alignment, averaging, and structural classification. Individual axonemal cross-sections were cropped and imported, and the software automatically centered and aligned microtubule doublets, the central pair, and dynein arm positions across images. Averaging reduced background noise and increased contrast, enabling clearer visualization of subtle defects such as inner dynein arm loss or central pair abnormalities.

**Basal Foot Orientation Analysis and Rose Plot Generation:** Basal foot orientation was quantified from TEM images using ImageJ/Fiji. Individual basal bodies with clearly identifiable basal feet were selected for analysis. For each basal body, a line was drawn from the basal body center to the tip of the basal foot using the Angle tool. ImageJ was configured to measure absolute angles with respect to the reference axis (*Analyze → Set Measurements → Angle = Absolute*). Angles were measured in degrees, rotating clockwise from the North reference. Values were exported to a .csv file using *Analyze → Measure*. Angle datasets were processed in either Python. Angles were grouped into consistent bin sizes (10°–20° depending on sample size) and plotted as rose diagrams. Each bin's radius represented the number of basal bodies within that orientation range. All plots maintained identical bin widths and radius scales for direct comparison across genotypes. Directional uniformity and mean direction differences were assessed using the Watson–Williams test, and circular variance was calculated for each sample.

**Quantification and Statistical Analysis**: All data were imported into GraphPad Prism for statistical analysis and graph generation. Comparisons between two groups were performed using a *t*-test with Welch's correction. Comparisons among more than two groups were conducted using one-way analysis of variance (ANOVA) followed by Tukey's post hoc test to determine statistical significance between groups, unless otherwise specified in the figure legends.

### SUPPLEMENTAL TABLES

#### Table E1. Human bronchial epithelial cell (HBEC) information

| | Lab Identity | Core Cell Demographics | |
|---|---|---|---|
| | | Sex | Age |
| CCDC40 mutant | L-65-PCD | Male | 41 |
| Wild-type Donor 1 | L-048-H | Male | 63 |
| Wild-type Donor 2 | L-085-H | Male | 63 |
| Wild-type Donor 3 | L-082-H | Female | 59 |
| Wild-type Donor 4 | L-083-H | Male | 63 |
| Wild-type Donor 5 | L-021-IPF | Male | 67* |
| Wild-type Donor 6 | L-004-H | Male | Unknown |

*Donor 5 is only included for reference in the supplemental information as was excluded from combined data due to being from an idiopathic pulmonary fibrosis patient (IPF) and thus not a wild-type pairing.

### TABLE E2. Primer pairs used for gene expression analysis

| Primer | Sequence |
|---|---|
| ODF2 F | GTG TCG CTC CTG GTT TCC AT |
| ODF2 R | TTC ATG GTT GGC TTC TGG CA |
| RPLP0 F | CCG TGA TGC CCA GGG AAG AC |
| RPLP0 R | GCA TCT GCT TGG AGC CCA CA |
| CCNO F | CCA GAG CTG CTA CGC CTT CC |
| CCNO R | GTC AGG CAC AGC GAC TCG AA |
| DNAi1 Ex5-6 F | TGG CAG TTC ACT ACA CCC AGG TT |
| DNAi1 Ex5-6 R | GCA GGC ACA TCT GTT TGA CTC CC |
| CCDC40 R | CCA GGT CTG AGG ACT CGA TGT |
| CCDC40 F | CCA GCA GTG GGC AGA TTG A |
| DNAH5 R | GGT CAT CCA GAG GCG GAA CG |
| DNAH5 F | CTG ATC CAC GGA CGC CAC TC |
| HPRT1 F | CAG CCC TGG CGT CGT GAT TA |
| HPRT1 R | TGT GAT GGC CTC CCA TCT CCT |
| MCIDAS R | GTT CGG CTG GCG AGT TCC TT |
| MCIDAS F | TGG CGG ACC AGA ACC AGA GA |
| TNFa R | GGG TTC GAG AAG ATG ATC TGA C |
| TNFa F | CAG GTT CTC TTC CTC TCA CAT AC |
| CC10 F | ACC ATG AAA CTC GCT GTC |
| CC10 R | TCA TAA CTG GAG GGT GTG TCC |
| FOXJ1 F | GTG CTT CAT CAA AGT GCC TCG |
| FOXJ1 R | GCC TCG GTA TTC ACC GTC AG |
| MUC5AC F | ACC AAT GCT CTG TAT CCT TCC C |
| MUC5AC R | TGG TGG ACG GAC AGT CAC T |

**TABLE E3. Antibodies used for IF Staining**

| Protein | Gene | Company/Catalogue # |
|---|---|---|
| Acetylated alpha tubulin | ATUB | Abcam ab24610 (1:500) mouse monoclonal |
| Radial Spoke Head Component 9 | RSPH9 | Sigma HPA031703 (1:500) rabbit polyclonal |
| Club Cell 10kDa | SCGB1A1 | R&D Systems MAB4218 (1:500) rat monoclonal |
| Mucin 5 AC | MUC5AC | Abcam ab198294 (1:250) rabbit monoclonal |

### SUPPLEMENTAL FIGURES

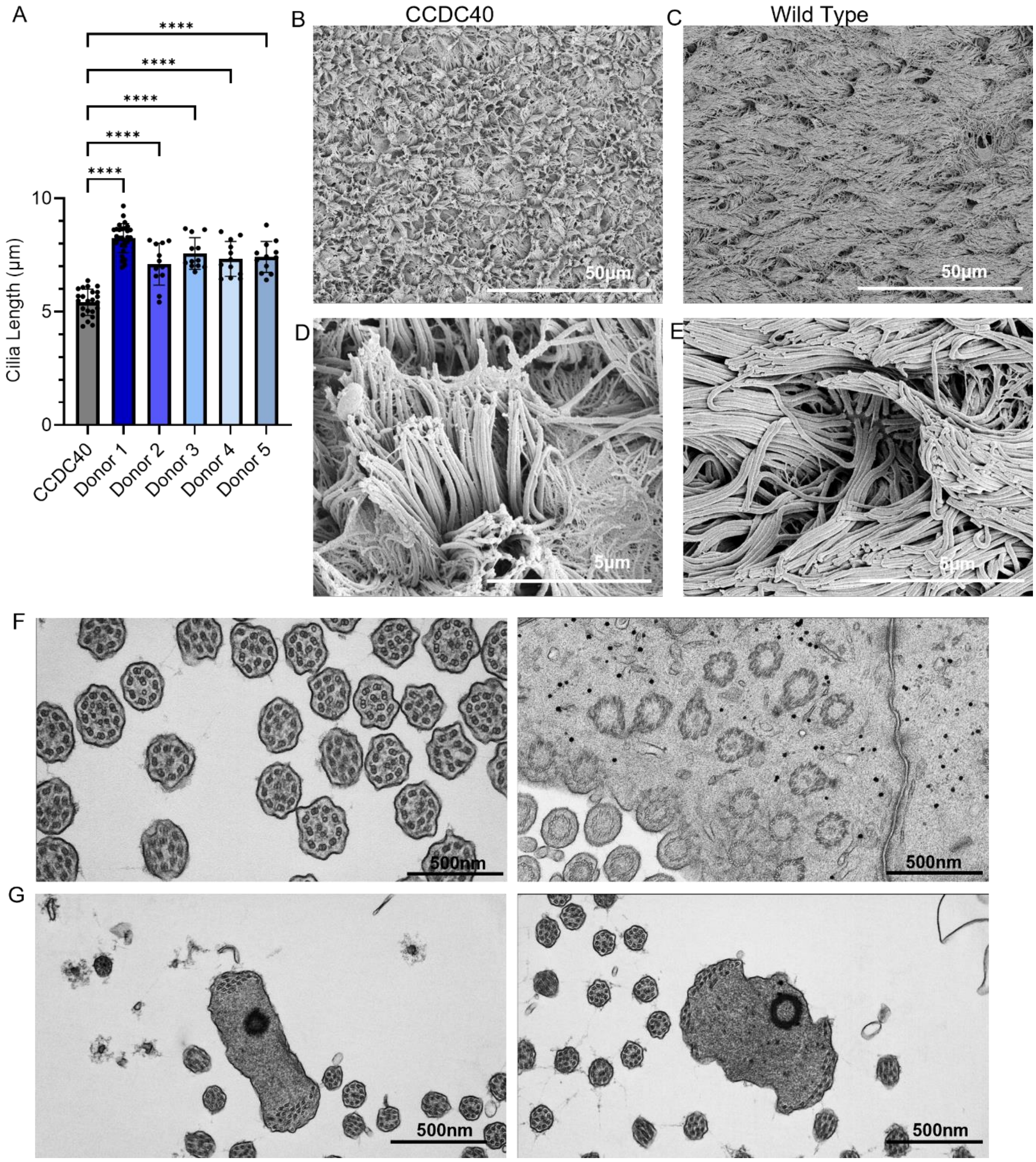


**Figure E1:** Supporting Figure 1 & 2: **CCDC40 mutant donor-derived ciliated cell have shorter, less uniform cilia**. **A**) Ciliary length per donor, each for represents an individual field of view from a minimum of three independent ALI differentiations. Data represents mean ± SEM, one way ANOVA ****$p<0.0001$. **B-E**) SEM images for CCDC40 mutant and Wild Type donors shown at two different magnifications. **F**) TEM images highlighting abnormal centriole replication and arrangements and misalignment of ciliary basal feet. **G**) TEM images highlighting unusual centriole-derived microtubule sets emerging at distant and opposing positions in enlarged structures.

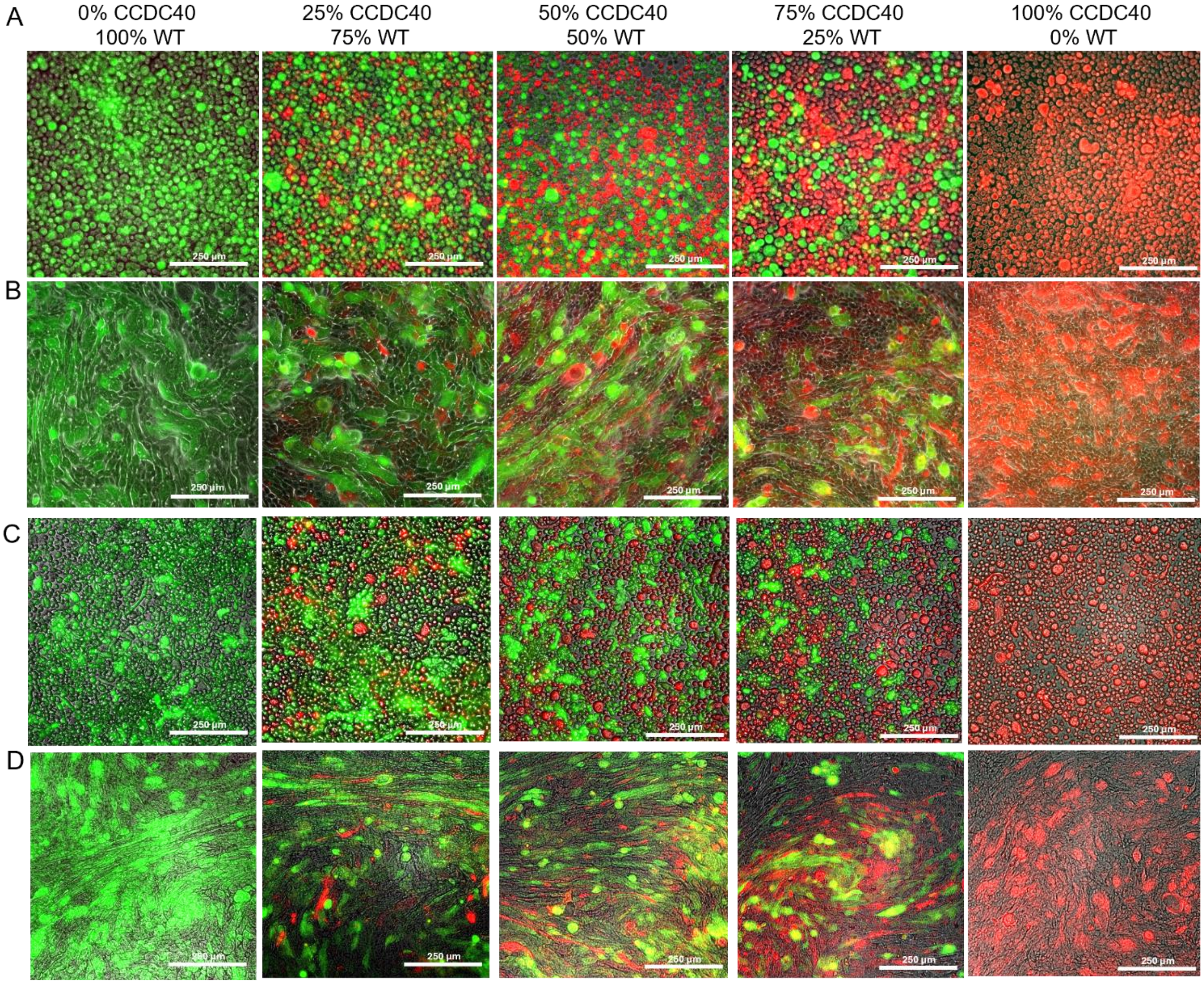


**Figure E2:** Supporting Figure 3: **Distribution of different ratios of mutant and wild-type donor HBEC remains similar during pre-ALI expansion**. Representative IF images of Cell Tracker Red labelled CCDC40 mutant HBEC and CellTracker Green labelled WT donor HBEC for an additional two donors. **A** and **C** are shown at day 0 of airlift and **B** and **D** are shown after 3 days of airlift across all plating ratios. Scale bars in all images are 250µm.

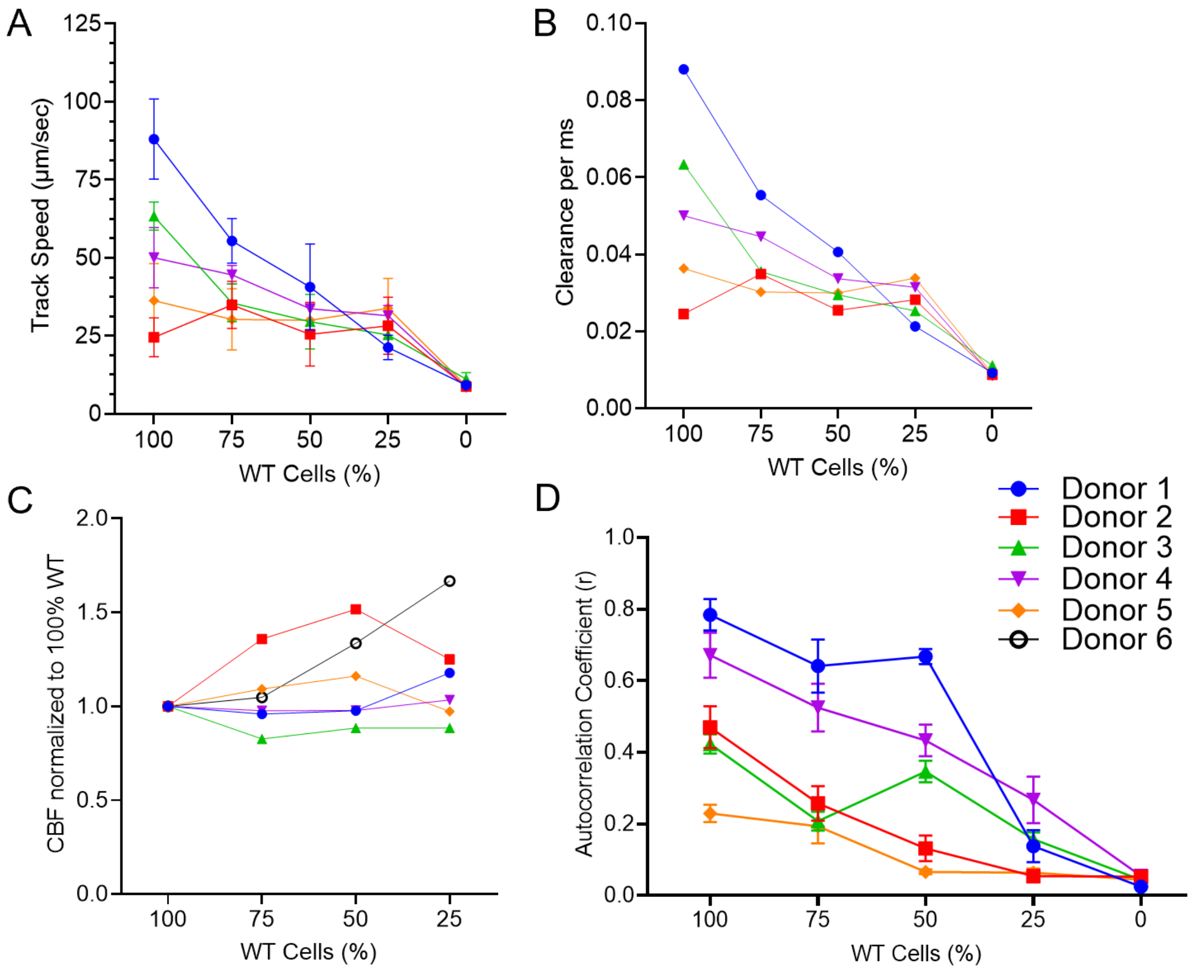


**Figure E3:** Supporting Figure 5: **Distribution of WT and CCDC40 cells regulates functional trajectories.** Normalized track speed (**A**) measured particle clearance speed from tracked fluorescent bead movement (**B**), CBF (**C**) and autocorrelation co-efficient (**D**) with lines representing each donor. Data represents average of technical replicates from 5-6 independent biological replicates for each experiment.
